# Identification of full-sibling families from natural single-tree ash progenies based on SSR markers and genome-wide SNPs

**DOI:** 10.1101/2023.07.18.549475

**Authors:** Melina Krautwurst, Franziska Past, Birgit Kersten, Ben Bubner, Niels A. Müller

## Abstract

Common ash, *Fraxinus excelsior*, is threatened by the invasive pathogen *Hymenoscyphus fraxineus*, which causes ash dieback. The pathogen is rapidly spreading throughout Europe with severe ecological and economic consequences. Multiple studies have presented evidence for the existence of a small fraction of genotypes with low susceptibility. Such genotypes can be targets for natural and artificial selection to conserve *F. excelsior* and associated ecosystems. To resolve the genetic architecture of variation in susceptibility it is necessary to analyze segregating populations. Here we employed about 1,000 individuals of each of four single-tree progenies from potentially tolerant mother trees to identify full-sibling (full-sib) families. To this end, we first genotyped all 4,000 individuals and the four mothers with eight SSR markers. We then used the program Colony to predict full-sibs without knowledge of the paternal genotypes. For each single-tree progeny, Colony predicted dozens of full-sib families, ranging from 3-165 individuals. In a next step, 910 individuals assigned to full-sib families with more than 30 individuals were subjected to high-resolution genotyping using over one million genome-wide SNPs which were identified with Illumina low-coverage resequencing. Using these SNP genotyping data in principal component analyses we were able to assign individuals to full-sib families with high confidence. Together the analyses revealed five large families with 80-212 individuals. These can be used to generate genetic linkage maps and to perform quantitative trait locus analyses for ash dieback susceptibility or other traits to *H. fraxineus* or other traits. The elucidation of the genetic basis of natural variation in ash may support breeding and conservation efforts and may contribute to more robust forest ecosystems.

## Introduction

Pests and diseases can cause severe diversity loss in woodlands, impacting ecological and economic aspects (Oliva et al. 2013; Hansen 1999; Johnson et al. 2015). In the early 1990s the necrotrophic ascomycete fungus *Hymenoscyphus fraxineus*, causing the ash dieback (ADB) disease, was first observed in Europe (Coker et al. 2019; Evans 2019). The fungus spread from its first distribution in Poland via wind-borne spores over most European ash populations (Kowalski 2006). The fungus can be traced back to Eastern Asia where it is associated with native *Fraxinus* species (McKinney et al. 2012; Husson et al. 2011; Zhao et al. 2013; Landolt et al. 2016). In Europe, the host of the pathogenic fungus is common ash (*Fraxinus excelsior*). Infected trees suffer from crown dieback and necrotic lesions. In the end, infection often leads to the death of the trees causing severe losses to European woodlands (Coker et al. 2019; Bakys et al. 2013).

Notably, *Fraxinus excelsior* exhibits natural variation in ADB susceptibility and several studies have shown that part of this variation is heritable with estimated heritabilities of 0.25 – 0.57 (Plumb et al. 2020; Enderle et al. 2015; Lobo et al. 2015; Lobo et al. 2014; McKinney et al. 2011; McKinney et al. 2014; McKinney et al. 2012; Muñoz et al. 2016; Pliura AL 2007; Pliura A et al. 2011; Harper et al. 2016). A number of different phenotypes are presumedly connected to ADB susceptibility. For example, the timing of bud burst and senescence may be important for variation in ADB although the details are still controversial (Nielsen et al. 2017; McKinney et al. 2011; Stener 2013; Pliura et al. 2016; Bakys et al. 2013).

Genotype-phenotype associations could reveal the genetic basis of variation in ADB and highlight candidate genes. Common ash is a wind-pollinated and wind-dispersed, polygamous subdioecious tree species. For genetic mapping or pedigree studies, selection of parents and artificial mating needs to be conducted. In natural tree populations it can be challenging to perform artificial mating or to identify the father to the naturally occurring seedlings, especially with a complex mating system as in *F. excelsior*. The method “breeding-without-breeding” (BwB) overcomes the need for artificial mating, by working with paternally unknown but maternally known material. Mothers can be selected based on their genotype or phenotype. Paired with DNA markers, it is possible to reconstruct pedigree structures with BwB and to use the identified full-sib families for quantitative trait locus analyses or the assessment of various breeding values (Lstibůrek et al. 2015; Lstibůrek et al. 2011). For pedigree prediction choosing a suitable downstream analysis for the sample set is important. Different molecular markers can be effective to assess kinships, such as simple sequence repeat (SSR) and single-nucleotide polymorphism (SNP) (Amom et al. 2020; Jiang et al. 2020; Zeng et al. 2023b).

SSR and SNP markers can be powerful tools in combination or separately (García et al. 2018; Zavinon et al. 2020; Capo-chichi et al. 2022; Zeng et al. 2023a). SNPs offer the opportunity to identify single base changes between individuals, are mostly biallelic, as well as the most abundant source of genetic polymorphism (Agarwal et al. 2008). With new sequencing technologies high numbers of SNPs can be reliably identified (Howe et al. 2020). SSRs are multi-allelic, highly polymorphic and currently the cheaper option for kinship assessment compared to SNPs, for which sequencing needs to be performed (Ramesh et al. 2020).

We therefore used eight SSR markers for genotyping of four single-tree progenies with 960 individuals each, which were collected from four potentially tolerant mother trees in Northern Germany. All four trees were closely monitored over several years. The SSR markers provide a low-cost high-throughput method for the prediction of potential full-sib families. Selected potential full-sibs were employed for high-resolution genotyping with low-coverage whole genome Illumina resequencing. Notably, only about half of the predicted families could be confirmed by the genome-wide SNP markers emphasizing the importance of considering different marker types for reliable identification of family structures. Nevertheless, our results demonstrate the overall feasibility of BwB in common ash. The validated families will be used for linkage mapping as part of the FraxForFuture project (Langer et al. 2022).

## Material and Methods

### Plant material

The four mother trees are distributed across the state of Mecklenburg-Western Pommerania in the north-east of Germany (Table 1). These trees were selected using an assessment scheme developed in a previous project (ResEsche). This scheme should ensure the vitality and silvicultural quality of the selected trees. The vitality criteria are assessed on the basis of foliage, shoot and trunk damage. The quality is recorded with parameters such as diameter at breast height (DBH), height and trunk shape. The selected mother trees all showed no or only few dieback symptoms in the crown area (no more than 10% crown defoliation, no more than 15% replacement shoot proportion). The upright-growth was of perfect or at least of normal quality (straight/upright growth, weak twisted growth, solid woodiness, etc.). Three of the four mother trees show no signs of ADB, only Eve-2 has a slight stem necrosis. Around 3000 seeds per tree were collected in 2018 as green seeds. All seeds were sown in a nursery bed within two weeks after harvesting. After germinating in Spring 2019, 960 seedlings per progeny were planted in 6×4 QuickPot plates (24 pots, 16 cm deep, HerkuPlast Kubern GmbH, Ering, Germany). They were first cultivated in a greenhouse and then transferred under a shading net in the nursery. In September 2019 plants were re-potted in 4×3 QuickPot plates (12 pots, 18 cm deep, HerkuPlast Kubern GmbH, Ering, Germany) and stayed in the nursery until planting. From 15^th^ to 17^th^ of April 2021 the seedlings were planted in a semi-randomized block design at a trial site near Schulzendorf (Brandenburg, Germany; Table 1). The area around the trial site is characterized by agriculture and small forests. Infected trees of *F. excelsior* with ash dieback symptoms were observed adjacent to the trial site.

**Table 1.**
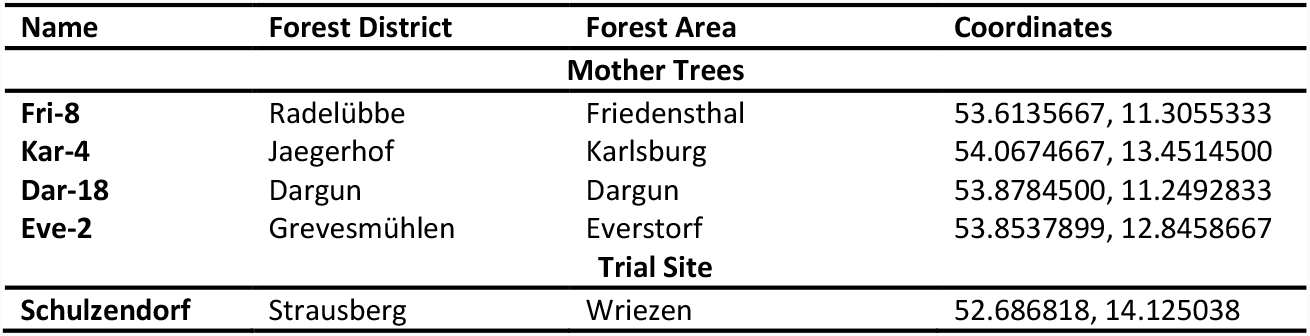
Location information of mother trees and trial site.

### Sampling, DNA extraction and SSR genotyping

Sampling for SSR genotyping was conducted in the nursery in late summer 2019 (Fri-8) and early summer 2020 (other progenies). The QuickPots were arranged in a 96-well-plate-like format. This arrangement allowed sampling in 96-well-plates without the need for time-consuming individual labeling. The 96-well-plates intended for DNA extraction were filled with two ceramic beads (1,4 mm Omni Beads, Omni International, Kennesaw; United States) per well using the customized Brendan bead dispenser (https://customlabinstitute.wordpress.com). The plate was cooled during the sampling process with a plate fitting ice pack. The sample, a 2 × 3 mm piece of the youngest, fully developed leaflet, was taken with forceps, which were cleaned with Ethanol (70 %) between each sample. After sampling, the plates were stored at -80 °C. Because of the large number of samples, a “quick and dirty”-method for DNA-extraction was chosen (Hu et al. 2014). The samples in the frozen sampling plates were homogenized (30 Hz; MM400, Retsch, Haan, Germany) for at least four minutes in precooled container. Depending on homogenization grade, additional homogenization was performed. After adding 200μl buffer (50 mM Tris, pH 8; 300 mM NaCl, 0,1 g/ml saccharose), the plates were again homogenized for another minute. After centrifuging the plate (5,889 x g; 5 min), the upper phase, containing the DNA, was directly used for polymerase chain reaction (PCR) without dilution or after diluting 1:10 on the same day. Plates with the “quick and dirty” DNA-extract were stored at -20 °C and, after another round of centrifugation, could be used for another PCR.

The PCR was performed with the Multiplex PCR Kit (Type-It Microsatellite master mix; Qiagen, Hilden, Germany). The SSR primer mix consisted of eight primers (see Table 2 and Table 3). With this SSR-multiplex, a touchdown procedure was performed (Table 4). The first progeny (Fri-8) was analyzed with F24 which was later replaced by F12, because F12 was more variable and more reliable.

**Table 2.** SSR-primers used for genotyping of ash seedlings in a multiplex assay. Primer are based on reference genome BATG0.5 (Sollars et al. 2017).

| primer | locus name | final concentration | label (color) | reference |
| --- | --- | --- | --- | --- |
| F12 | FEMSATL12C | 0.2 µM | DY-751 (blue) | adapted FEMSATL12 (Lefort et al. 1999) |
| F23 | Contig184_179564_179439 | 0.2 µM | DY-682 (green) | (Sollars et al. 2017) |
| F25 | Contig1992_131610_131822 | 0.2 µM | DY-682 (green) | (Sollars et al. 2017) |
| F26 | Contig918_100265_100431 | 0.2 µM | DY-751 (blue) | (Sollars et al. 2017) |
| F27 | Contig5418_11372_11513 | 0.2 µM | Cy5 (black) | (Sollars et al. 2017) |
| F28 | Contig3670_10994_10705 | 0.4 µM | DY-682 (green) | (Sollars et al. 2017) |
| F29 | Contig4738_19258_19142 | 0.2 µM | DY-751 (blue) | (Sollars et al. 2017) |
| F30 | Contig344_81629_81375 | 0.2 µM | Cy5 (black) | (Sollars et al. 2017) |
| F24 | Contig5748_109580_109480 | 0.1 µM | DY-751 (blue) | (Sollars et al. 2017) |

**Table 3.**
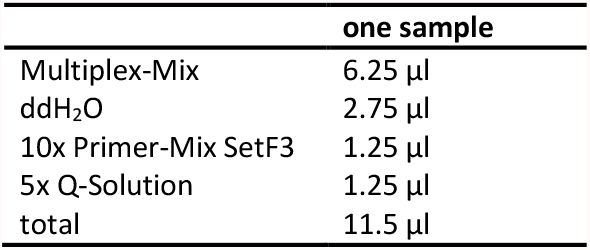
PCR master mix.

|  | one sample |
| --- | --- |
| Multiplex-Mix | 6.25 µl |
| ddH<sub>2</sub>O | 2.75 µl |
| 10x Primer-Mix SetF3 | 1.25 µl |
| 5x Q-Solution | 1.25 µl |
| total | 11.5 µl |

**Table 4.**
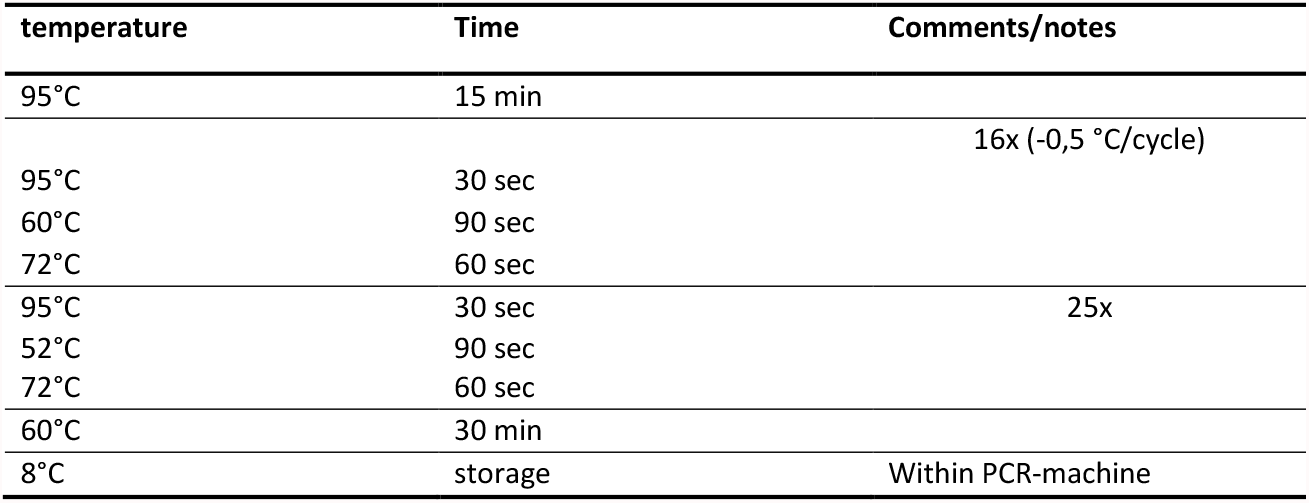
Polymerase chain reaction program.

**Table 5:** Summary of all full-sibling families identified by SSR and SNP marker.

| Progeny | Abbreviation | SSR-name | Size | SNP-name | Size |
| --- | --- | --- | --- | --- | --- |
| Friedrichsthal | Fri-8 | F10 | 81 | Cluster 1 | 212 |
|  |  | F17 | 90 | Cluster 2 | 26 |
|  |  | F22 | 63 |  |  |
| Karlsburg | Kar-4 | F29 | 112 | Cluster 1 | 164 |
|  |  | F05 | 36 | Cluster 2 | 25 |
|  |  | F21 | 30 |  |  |
|  |  | F41 | 33 |  |  |
| Dargun | Dar-18 | F20 | 165 | Cluster 2 | 138 |
|  |  | F08 | 85 | Cluster 1 | 73 |
|  |  | F30 | 39 | Cluster 3 | 35 |
| Everstorf | Eve-2 | F19 | 124 | Cluster1 | 116 |
|  |  | F02 | 31 |  |  |

The PCR products were analyzed with capillary electrophoresis (GenomeLab™ GeXP, Beckman Coulter) and the corresponding chemical kit (SCIEX). The peak scoring was done with the provided software (GenomeLab) to obtain a list of the alleles (Table S1-S4).

### Colony analysis

In order to determine genetic sample relationships, the SSR-genotyping output was transferred to the COLONY Software (Jones and Wang 2010, Version: 2.0.6.5/2.0.6.6). In addition to the input data, input parameters for the analysis of all progenies had to be defined (see Supplement: Supplemental materials on colony parameters for family estimations with SSR markers). The software works with likelihood(s). This can lead to different outputs, if the analysis is repeated. To ensure reliability of the results, the software was run twice. The two runs were compared and the individuals that were observed in both runs were chosen.

### DNA extraction and Illumina low-coverage resequencing

The first batch of 167 samples with young leaf material was collected in September 2020, immediately placed on ice in 1.5 ml Eppendorf safe-lock tubes (Eppendorf, Wesslingen, Germany). In addition, leaves from the four mother tree clones (Table 1) were collected. All samples were stored at -80°C until DNA extraction. Frozen samples were homogenized using pestle and mortar in liquid nitrogen. All following steps were conducted following the DNA extraction protocol by (Bruegmann et al. 2022, adapted with DL_Dithiothreitol). The second batch of 751 samples was collected in June 2021. For the 751 samples we used the MagMAX™ Plant DNA Isolation Kit (Thermo Fisher Scientific, Germany) following purification using the KingFisher™ Apex (Thermo Fisher Scientific, Germany) with a 96 deep-well head. DNA sample QC and library preparation for sequencing were performed by Novogene (UK) Ltd. (Cambridge, UK) for both sample batches. In the first batch, 163 of 172 samples passed the quality control, including the four mother tree samples. In the second batch, 747 of 751 samples passed the quality control. Sequencing data (2 × 150 bp reads) were generated on the Novaseq 6000 platform for both batches. The first batch was sequenced to an average sequencing depth of 10.8X and the second batch to 11.3X according to the ash reference genome (Sollars et al. 2017). Both batches together comprise 906 samples plus the four mother trees.

### Mapping and variant calling

For both batches, sequencing data were mapped against the common ash reference genome (Sollars et al. 2017) using bwa-mem (version bwa-0.7.17.tar.bz2 (Li and Durbin 2009)). Grouping of the reads and duplicates were marked using Picard tool’s (v2.26.2) (http://broadinstitute.github.io/picard/). Joint variant calling was performed for batch one with GATK (version 4.2.3.0), following the best practices for germline short variant discovery wherever possible (Poplin et al. 2017). For the second batch the variant calling was performed by Novogene using Sentieon (Aldana and Freed 2022). For generating gVCFs the ‘HaplotypeCaller‘ from GATK was used. After combining the gVCFs with GATK’s ‘GenomicsDBImport’, the ‘GenotypeGVFs’ tool was used for the joint genotyping. For batch one GATK version 4.2.3.0 and for batch two GATK version 4.0.5.1 were used (van der Auwera and O’Connor 2020).

### Variant filtering

The two sample sets were filtered separately, with the same filtering options. For hard filtering we mostly followed the documentation on ‘Hard-filtering germline short variants’ on the GATK website. We filtered indels and SNPs separately. We removed variants based on strand bias (FisherStrand ‘FS’ > 60 & StrandOddsRatio ‘SOR’ > 3) and mapping quality (RMSMappingQuality ‘MQ’ < 40, MappingQualityRankSumTest ‘MQRankSum’ < -1). Based on the distribution of the variant confidence score QualByDepth ‘QD’ we chose a more stringent cutoff of QD > 10, to remove any low-confidence variants. Filtering was performed with bcftools v1.7 (Li 2011). We then extracted the variant sequencing depth values ‘DP’ and minor allele frequency ‘frq2‘ using vcftools v0.1.15 (Danecek et al. 2011). To visualize the DP and choose the parameters we used R (R Core Team 2022). Minimal-mean DP was 5.2 and max-mean DP 10.3. Non-biallelic SNPs were excluded and SNPs with more than 10 % missing data were removed. Filtering resulted in a set of 14.42 million SNPs for the first batch and 11.78 million SNPs for the second batch. Before the resulting VCF files (Variant Call Format) could be merged, an intersect was calculated using the ‘isec’ function of bcftools (Danecek et al. 2021) to identify common SNPs in both VCFs files. Then, the ‘merge’ function was used to create one multi-sample file. Further, the minor allele frequency was filtered 0.08-0.5 with PLINK v1.9 (Purcell et al. 2007).The merged file of both batches comprised 5.87 million SNPs for 910 individuals.

We used PLINK for principal component analyses (PCA) to determine family-clusters in the SNP datasets. Further, we used the R package ‘SNPRelate’ (Zheng et al. 2012) for the genome-wide identity-by-state analysis to create a dendrogram based on the SNP data.

### Data visualization

The results were further analyzed and visualized using the R packages ‘ggplot2’ (Wickham 2016), ‘VCFR’ (Knaus and Grünwald 2017), ‘MASS’ (Venables and Ripley 2003) and ‘dendextend’ (Galili 2015).

## Results

A total of 960 samples per single-tree progeny of each of four mother tree, potentially tolerant against ash dieback, were collected and analyzed with eight SSR markers (Table S1-S4). In the end, 3.476 of 3.840 (90.5 %) samples could be successfully genotyped. The program Colony predicted between 151 and 179 full-sib families per progeny (Figure 1 and Figure 2) including families with 1-165 members. In total, 910 individuals, which were assigned to the nine largest predicted full-sib families, were selected for high-resolution genotyping using Illumina whole-genome resequencing. Only families with at least 30 individuals were chosen. With a sequencing depth of 10X we were able to identify a total 5.87 million SNPs after read mapping to the *F. excelsior* reference genome (Sollars et al. 2017) and SNP filtering. Employing only SNPs that are largely independent (r2 < 0.2) within each progeny, we were able to reliably assign full-sib families. The final file included six samples that were sequenced and analyzed in batch 1 and batch 2. With these ‘twins’ we were able to compare the two downstream analyses. The ‘twins’ are represented in the Fri-8 dendrogram (Figure 2), which shows the same results for both downstream analysis.

**Figure 1.**
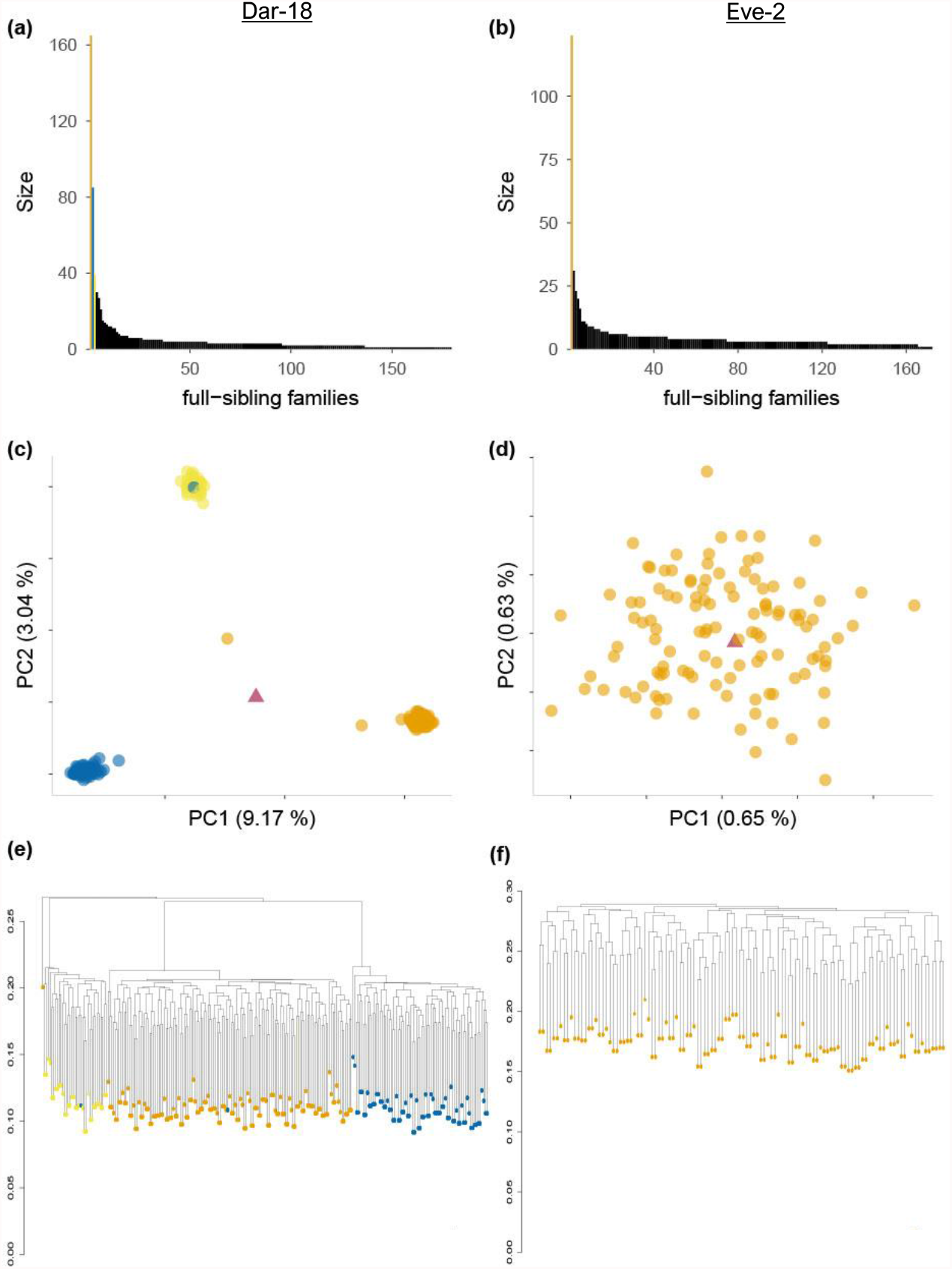
Comparison of SSR and SNP full-sibling identification from the mother trees Dar-18 and Eve-2. The barplots (a and b) show all progenies that were analyzed with the SSR markers and assigned to full-sibling families (range from 1 – 165 siblings). Each bar represents one predicted full-sibling family. Selected families with more than 30 individuals are indicated by colors, that is three families for Dar-18 (a) and one family for Eve-2 (b). The principal component analyses (c and d) show the SNP marker results. The color scheme of the dots represents the results of the SSR markers. The red triangle represents the mother tree. Panel (e) and (f) show the results of genome-wide identity-by-state analyses using the SNP markers results in dendrograms. The dots represent the SSR results and the dendrogram clustering represents the SNP markers.

**Figure 2:**
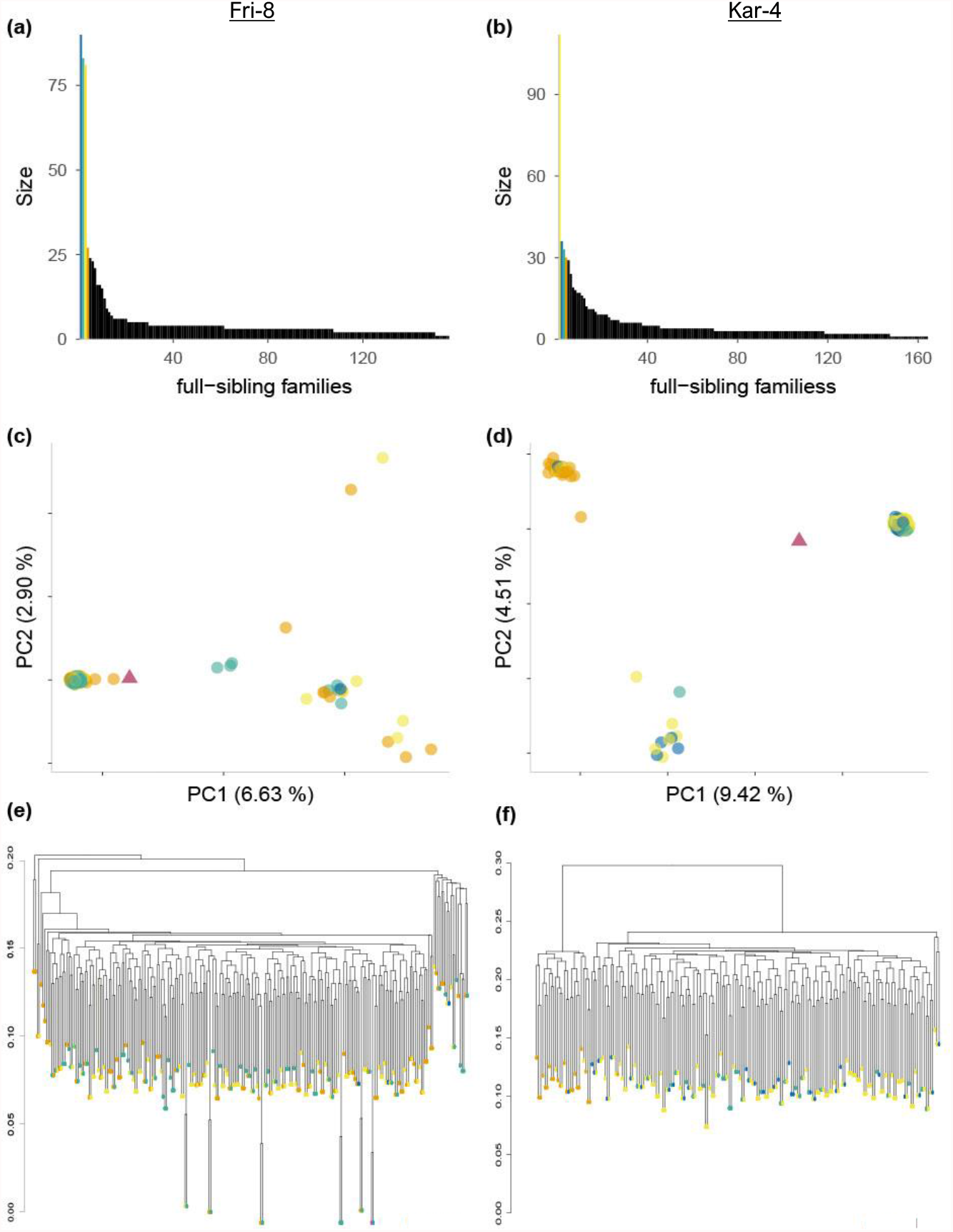
Comparison of SSR and SNP full-sibling identification from the mother trees Fri-8 and Kar-4. The barplots (a and b) show all progenies that were analyzed with the SSR markers and assigned to full-sibling families (range from 1 – 112 siblings). Each bar represents one predicted full-sibling family. Selected families with more than 30 individuals are indicated by colors, that is four families for Fri-8 (a) and four families for Kar-4 (b). The principal component analyses (c and d) show SNP marker results. The color scheme of the dots represents the results of the SSR markers. The red triangle represents the mother tree. Panel (e) and (f) show the results of genome-wide identity-by-state analysis using the SNP markers in dendrograms. The dots represent the SSR results and the dendrogram clustering represents the SNP markers.

For the two single-tree progenies from Dar-18 and Eve-2, the family structure predicted by Colony using the eight SSR markers was largely consistent with the SNP data analyses (Figure. 1). Only a few outliers were detected. Accordingly, family sizes were close to the predictions with the largest full-sib family from Dar-18 comprising 138 individuals (Table 1). For Eve-2, only a single dominant pollen donor gave rise to a single large full-sib family, of which 116 individuals could be confirmed with the SNP markers. Whereas Dar-18 and Eve-2 demonstrate the general feasibility of performing breeding-without-breeding in ash with the eight described SSR markers, the other two single-tree progenies from Kar-4 and Fri-8 showed discrepancies between the SRR and SNP marker classifications (Figure 2). Especially Fri-8, which was predicted to be composed of four full-sib families using SSR markers, turned out to represent a single large family of 212 individuals based on SNP markers. Additional individuals appear as outliers representing other pollen donors. Finally, for Kar-4, our SNP marker analysis shows two relatively large families (164 and 25 individuals) instead of four. While breeding-without-breeding appears to work well for common ash in principle, our results also highlight the importance of validating family predictions with different types of markers, or higher numbers of informative SSR markers.

## Discussion

For genotype-phenotype association studies it is important to maximize the statistical power by employing large mapping populations with a high number of individuals. In BwB approaches the reliable determination of full-sib families is critical. Although whole-genome resequencing and genome-wide SNP detection provide robust data for this purpose, sequencing costs can still be prohibitive with thousands of samples, such as the 4,000 individuals in our study. The preselection of individuals prior to SNP genotyping can thus be important, and SSR markers provide the required simplicity and relatively low costs. This is reflected by ongoing research with SSR markers, identification of polymorphic SSRs being only one example (Mishra et al. 2023; Guichoux et al. 2011).

Our results show, that BwB implementing a two-step genotyping with SSR and SNP markers is an effective way to achieve the identification of full-sib families in ash. While the SSRs can be used to do a preselection of samples based on full-sib prediction and avoid sequencing of less relevant individuals, the genome-wide SNP data can be employed for full-sib validation and construction of genetic maps. Despite some discrepancies between SSR and SNP marker classification, the preselection of individuals allowed us to identify relatively large full-sib families for deeper analysis by SNP genotyping. Similar studies have shown comparable results (van Inghelandt et al. 2010). In a study from et al Zavinon (2020) family identification in Beninese pigeon pea (*Cajanus cajan* (L.)) populations was conducted with 30 informative SSR loci and 794 genotype by sequencing (GBS) derived SNPs. The results with both marker sets were similar, but the PCA based on SNP markers showed the more accurate results. For genotype-phenotype association studies, crossing and back-crossing generations are essential. For tree species with long generation times, this can be challenging. With the BwB technique that we used, we were able to generate F1 generations and full-sib families from four different tolerant mother trees. The combination of SSR and SNP markers enabled successful identification of families that are a valuable resource to perform quantitative trait locus analyses for susceptibility to *H. fraxineus* in future studies.

## Acknowledgements

We thank our technical assistants Viktoria Blunk, Annika Eikhof, Katrin Groppe, Marlies Karaus, Heidrun Mattauch, and Regina Zimmermann for their help with laboratory and field work. We thank members of the FraxForFuture project and the Thünen Institute of Forest Genetics for experimental advice and discussion on the project and data analysis. The FraxForFuture research network is funded by the German Federal Ministry of Food and Agriculture and the German Federal Ministry for the Environment, Nature Conservation and Nuclear Safety. The projects of the sub-networks are funded by the Waldklimafonds and the Thünen Institute. The project executing agency is the Fachagentur für Nachwachsende Rohstoffe e.V. (FNR).

## Data availability

All genomic variants have been deposited in the European Variant Archive (EVA) under the accession number ‘PRJEB64325’.

## Supplementary Materials

### Supplemental materials on colony parameters for family estimations with SSR marker

For the analytical process with the program colony following parameters were chosen: Mating system: male polygam and female polygam, with inbreeding, without clones, dioecious and diploid, length of run: medium, analysis method: full likelihood, likelihood precision: high, run specification: default and no sibship prior.

Table S1: Allele report of offspring genotypes of Fri-8 of the program colony.

Table S2: Allele report of offspring genotypes of Eve-2 of the program colony.

Table S3: Allele report of offspring genotypes of Kar -4 of the program colony.

Table S4: Allele report of offspring genotypes of Dar-18 of the program colony.

## References

Agarwal, Milee; Shrivastava, Neeta; Padh, Harish (2008): Advances in molecular marker techniques and their applications in plant sciences. In Plant Cell Rep 27 (4), pp. 617–631. DOI: 10.1007/s00299-008-0507-z.

Aldana, Rafael; Freed, Donald (2022): Data Processing and Germline Variant Calling with the Sentieon Pipeline. In Methods in molecular biology (Clifton, N.J.) 2493, pp. 1–19. DOI: 10.1007/978-1-0716-2293-3_1.

Amom, Thoungamba; Tikendra, Leimapokpam; Apana, Nandeibam; Goutam, Moirangthem; Sonia, Paonam; Koijam, Arunkumar Singh et al. (2020): Efficiency of RAPD, ISSR, iPBS, SCoT and phytochemical markers in the genetic relationship study of five native and economical important bamboos of North-East India. In Phytochemistry 174, p. 112330. DOI: 10.1016/j.phytochem.2020.112330.

Bakys, R., Vasaitis, R., Skovsgaard, J. P. (2013): Patterns and severity of crown dieback in young even-aged stands of european ash (Fraxinus excelsior L.) in relation to stand density, bud flushing phenotype, and season. In Plant Protect. Sci. 49 (No. 3), pp. 120–126. DOI: 10.17221/70/2012-PPS.

Bruegmann, Tobias; Fladung, Matthias; Schroeder, Hilke (2022): Flexible DNA isolation procedure for different tree species as a convenient lab routine. In Silvae Genetica 71 (1), pp. 20–30. DOI: 10.2478/sg-2022-0003.

Capo-chichi, Ludovic J. A., Elakhdar, Ammar; Kubo, Takahiko; Nyachiro, Joseph; Juskiw, Patricia; Capettini, Flavio et al. (2022): Genetic diversity and population structure assessment of Western Canadian barley cooperative trials. In Frontiers in plant science 13, p. 1006719. DOI: 10.3389/fpls.2022.1006719.

Coker, Tim L. R., Rozsypálek, Jiří; Edwards, Anne; Harwood, Tony P., Butfoy, Louise; Buggs, Richard J. (2019): Estimating mortality rates of European ash (Fraxinus excelsior) under the ash dieback (Hymenoscyphus fraxineus) epidemic. In Plants, People, Planet 1 (1), pp. 48–58. DOI: 10.1002/ppp3.11.

Danecek, Petr; Auton, Adam; Abecasis, Goncalo; Albers, Cornelis A., Banks, Eric; DePristo, Mark A. et al. (2011): The variant call format and VCFtools. In Bioinformatics (Oxford, England) 27 (15), pp. 2156–2158. DOI: 10.1093/bioinformatics/btr330.

Danecek, Petr; Bonfield, James K., Liddle, Jennifer; Marshall, John; Ohan, Valeriu; Pollard, Martin O. et al. (2021): Twelve years of SAMtools and BCFtools. In GigaScience 10 (2). DOI: 10.1093/gigascience/giab008.

Enderle, Rasmus; Nakou, Aikaterini; Thomas, Kristina; Metzler, Berthold (2015): Susceptibility of autochthonous German Fraxinus excelsior clones to Hymenoscyphus pseudoalbidus is genetically determined. In Annals of Forest Science 72 (2), pp. 183–193. DOI: 10.1007/s13595-014-0413-1.

Evans, Matthew R. (2019): Will natural resistance result in populations of ash trees remaining in British woodlands after a century of ash dieback disease? In Royal Society open science 6 (8), p. 190908. DOI: 10.1098/rsos.190908.

Galili, Tal (2015): dendextend: an R package for visualizing, adjusting and comparing trees of hierarchical clustering. In Bioinformatics (Oxford, England) 31 (22), pp. 3718–3720. DOI: 10.1093/bioinformatics/btv428.

García, Cristina; Guichoux, Erwan; Hampe, Arndt (2018): A comparative analysis between SNPs and SSRs to investigate genetic variation in a juniper species (Juniperus phoenicea ssp. turbinata). In Tree Genetics & Genomes 14 (6), pp. 1–9. DOI: 10.1007/s11295-018-1301-x.

Guichoux, E., Lagache, L., Wagner, S., Chaumeil, P., LÉger, P., Lepais, O. et al. (2011): Current trends in microsatellite genotyping. In Molecular ecology resources 11 (4), pp. 591–611. DOI: 10.1111/j.1755-0998.2011.03014.x.

Hansen, Everett M. (1999): Disease and diversity in forest ecosystems. In Austral. Plant Pathol. 28 (4), p. 313. DOI: 10.1071/AP99050.

Harper, Andrea L., McKinney, Lea Vig; Nielsen, Lene Rostgaard; Havlickova, Lenka; Li, Yi; Trick, Martin et al. (2016): Molecular markers for tolerance of European ash (Fraxinus excelsior) to dieback disease identified using Associative Transcriptomics. In Sci Rep 6 (1), p. 19335. DOI: 10.1038/srep19335.

Howe, Glenn T., Jayawickrama, Keith; Kolpak, Scott E., Kling, Jennifer; Trappe, Matt; Hipkins, Valerie et al. (2020): An Axiom SNP genotyping array for Douglas-fir. In BMC Genomics 21 (1), p. 9. DOI: 10.1186/s12864-019-6383-9.

Hu, Jin-Yong; Zhou, Yue; He, Fei; Dong, Xue; Liu, Liang-Yu; Coupland, George et al. (2014): miR824-Regulated AGAMOUS-LIKE16 Contributes to Flowering Time Repression in Arabidopsis. In The Plant cell 26 (5), pp. 2024–2037. DOI: 10.1105/tpc.114.124685.

Husson, Claude; Scala, Bruno; Caël, Olivier; Frey, Pascal; Feau, Nicolas; Ioos, Renaud; Marçais, Benoît (2011): Chalara fraxinea is an invasive pathogen in France. In Eur J Plant Pathol 130 (3), pp. 311–324. DOI: 10.1007/s10658-011-9755-9.

Jiang, Kaibin; Xie, Hui; Liu, Tianyi; Liu, Chunxin; Huang, Shaowei (2020): Genetic diversity and population structure in Castanopsis fissa revealed by analyses of sequence-related amplified polymorphism (SRAP) markers. In Tree Genetics & Genomes 16 (4), pp. 1–10. DOI: 10.1007/s11295-020-01442-2.

Johnson, Pieter T. J., Ostfeld, Richard S., Keesing, Felicia (2015): Frontiers in research on biodiversity and disease. In Ecology letters 18 (10), pp. 1119–1133. DOI: 10.1111/ele.12479.

Jones, Owen R., Wang, Jinliang (2010): COLONY: a program for parentage and sibship inference from multilocus genotype data. In Molecular ecology resources 10 (3), pp. 551–555.

Knaus, Brian J., Grünwald, Niklaus J. (2017): vcfr: a package to manipulate and visualize variant call format data in R. In Molecular ecology resources 17 (1), pp. 44–53. DOI: 10.1111/1755-0998.12549.

Kowalski, T. (2006): Chalara fraxinea sp. nov. associated with dieback of ash (Fraxinus excelsior) in Poland. In For. Path. 36 (4), pp. 264–270. DOI: 10.1111/j.1439-0329.2006.00453.x.

Landolt, J., Gross, A., Holdenrieder, O., Pautasso, M. (2016): Ash dieback due to Hymenoscyphus fraxineus : what can be learnt from evolutionary ecology? In Plant Pathol 65 (7), pp. 1056–1070. DOI: 10.1111/ppa.12539.

Langer, Gitta Jutta; Fuchs, Sebastian; Osewold, Johannes; Peters, Sandra; Schrewe, Falk; Ridley, Maia et al. (2022): FraxForFuture—research on European ash dieback in Germany. In J Plant Dis Prot, pp. 1–11. DOI: 10.1007/s41348-022-00670-z.

Lefort, F., Brachet, S., Frascaria-Lacoste, N., Edwards, K. J., Douglas, G. C. (1999): Identification and characterization of microsatellite loci in ash (Fraxinus excelsior L.) and their conservation in the olive family (Oleaceae). In Mol Ecol 8 (6), pp. 1088–1089. DOI: 10.1046/j.1365-294X.1999.00655_8.x.

Li, Heng (2011): A statistical framework for SNP calling, mutation discovery, association mapping and population genetical parameter estimation from sequencing data. In Bioinformatics (Oxford, England) 27 (21), pp. 2987–2993. DOI: 10.1093/bioinformatics/btr509.

Li, Heng; Durbin, Richard (2009): Fast and accurate short read alignment with Burrows-Wheeler transform. In Bioinformatics (Oxford, England) 25 (14), pp. 1754–1760. DOI: 10.1093/bioinformatics/btp324.

Lobo, A., McKinney, L. V., Hansen, J. K., Kjaer, E. D., Nielsen, L. R. (2015): Genetic variation in dieback resistance in Fraxinus excelsior confirmed by progeny inoculation assay. In For. Path. 45 (5), pp. 379–387. DOI: 10.1111/efp.12179.

Lobo, Albin; Hansen, Jon Kehlet; McKinney, Lea Vig; Nielsen, Lene Rostgaard; Kjær, Erik Dahl (2014): Genetic variation in dieback resistance: growth and survival of Fraxinus excelsior under the influence of Hymenoscyphus pseudoalbidus. In Scandinavian Journal of Forest Research 29 (6), pp. 519–526. DOI: 10.1080/02827581.2014.950603.

Lstibůrek, Milan; Hodge, Gary R., Lachout, Petr (2015): Uncovering genetic information from commercial forest plantations—making up for lost time using “Breeding without Breeding”. In Tree Genetics & Genomes 11 (3), pp. 1–12. DOI: 10.1007/s11295-015-0881-y.

Lstibůrek, Milan; Ivanková, Kristýna; Kadlec, Jan; Kobliha, Jaroslav; Klápště, Jaroslav; El-Kassaby, Yousry A. (2011): Breeding without breeding: minimum fingerprinting effort with respect to the effective population size. In Tree Genetics & Genomes 7 (5), pp. 1069–1078. DOI: 10.1007/s11295-011-0395-1.

McKinney, L. V., Nielsen, L. R., Collinge, D. B., Thomsen, I. M., Hansen, J. K., Kjaer, E. D. (2014): The ash dieback crisis: genetic variation in resistance can prove a long-term solution. In Plant Pathol 63 (3), pp. 485–499. DOI: 10.1111/ppa.12196.

McKinney, L. V., Nielsen, L. R., Hansen, J. K., Kjær, E. D. (2011): Presence of natural genetic resistance in Fraxinus excelsior (Oleraceae) to Chalara fraxinea (Ascomycota): an emerging infectious disease. In Heredity 106 (5), pp. 788–797. DOI: 10.1038/hdy.2010.119.

McKinney, L. V., Thomsen, I. M., Kjaer, E. D., Nielsen, L. R. (2012): Genetic resistance to Hymenoscyphus pseudoalbidus limits fungal growth and symptom occurrence in Fraxinus excelsior. In For. Path. 42 (1), pp. 69–74. DOI: 10.1111/j.1439-0329.2011.00725.x.

Mishra, Garima; Meena, Rajendra K., Kant, Rama; Pandey, Shailesh; Ginwal, Harish S., Bhandari, Maneesh S. (2023): Genome-wide characterization leading to simple sequence repeat (SSR) markers development in Shorea robusta. In Funct Integr Genomics 23 (1), pp. 1–16. DOI: 10.1007/s10142-023-00975-8.

Muñoz, Facundo; Marçais, Benoît; Dufour, Jean; Dowkiw, Arnaud (2016): Rising Out of the Ashes: Additive Genetic Variation for Crown and Collar Resistance to Hymenoscyphus fraxineus in Fraxinus excelsior. In Phytopathology 106 (12), pp. 1535–1543. DOI: 10.1094/PHYTO-11-15-0284-R.

Nielsen, L. R., McKinney, L. V., Kjær, E. D. (2017): Host phenological stage potentially affects dieback severity after Hymenoscyphus fraxineus infection in Fraxinus excelsior seedlings. In Baltic Forestry 23 (1), pp. 229–232. Available online at https://www.cabdirect.org/cabdirect/abstract/20173295909.

Oliva, J., Boberg, J. B., Hopkins, A. J. M., Stenlid, J. (2013): Concepts of epidemiology of forest diseases. In Paolo Gonthier, Giovanni Nicolotti, Luana Giordano (Eds.): Infectious forest diseases. Oxfordshire, England, Boston, Massachusetts: CABI, pp. 1–28.

Pliura, A., Lygis, V., Marčiulyniene, D., Suchockas, V., Bakys, R. (2016): Genetic variation of Fraxinus excelsior half-sib families in response to ash dieback disease following simulated spring frost and summer drought treatments. In iForest 9 (1), Article 1514, pp. 12–22. DOI: 10.3832/ifor1514-008.

Pliura A; Lygis V; Suchockas V; Bartkevicius E. (2011): Performance of twenty four European Fraxinus excelsior populations in three Lithuanian progeny trials with a special emphasis on resistance to Chalara … (17.1), pp. 17–34.

Pliura AL, Baliuckas V. I. (2007): Genetic variation in adaptive traits of progenies of Lithuanian and western European populations of Fraxinus excelsior L. In Baltic Forestry (13(1)), pp. 28–38. Available online at https://www.balticforestry.mi.lt/bf/pdf_articles/2007-13[1]/28_38%20pliura%20&%20baliuckas.pdf.

Plumb, William J., Coker, Timothy L. R., Stocks, Jonathan J., Woodcock, Paul; Quine, Christopher P., Nemesio-Gorriz, Miguel et al. (2020): The viability of a breeding programme for ash in the British Isles in the face of ash dieback. In Plants, People, Planet 2 (1), pp. 29–40. DOI: 10.1002/ppp3.10060.

Poplin, Ryan; Ruano-Rubio, Valentin; DePristo, Mark A., Fennell, Tim J., Carneiro, Mauricio O., van der Auwera, Geraldine A. et al. (2017): Scaling accurate genetic variant discovery to tens of thousands of samples.

Purcell, Shaun; Neale, Benjamin; Todd-Brown, Kathe; Thomas, Lori; Ferreira, Manuel A. R., Bender, David et al. (2007): PLINK: a tool set for whole-genome association and population-based linkage analyses. In The American Journal of Human Genetics 81 (3), pp. 559–575. DOI: 10.1086/519795.

R Core Team (2022): R: A language and environment for statistical computing. R Foundation for. Vienna, Austria. Available online at https://www.R-project.org/.

Ramesh, Palakurthi; Mallikarjuna, Gunti; Sameena, Shaik; Kumar, Anand; Gurulakshmi, Kola; Reddy, Vigneswara et al. (2020): Advancements in molecular marker technologies and their applications in diversity studies. In J Biosci 45 (1), pp. 1–15. DOI: 10.1007/s12038-020-00089-4.

Sollars, Elizabeth S. A., Harper, Andrea L., Kelly, Laura J., Sambles, Christine M., Ramirez-Gonzalez, Ricardo H., Swarbreck, David et al. (2017): Genome sequence and genetic diversity of European ash trees. In Nature 541 (7636), pp. 212–216. DOI: 10.1038/nature20786.

Stener, Lars-Göran (2013): Clonal differences in susceptibility to the dieback of Fraxinus excelsior in southern Sweden. In Scandinavian Journal of Forest Research 28 (3), pp. 205–216. DOI: 10.1080/02827581.2012.735699.

van der Auwera, G. A., O’Connor, B. D. (2020): Genomics in the Cloud: Using Docker, GATK, and WDL in Terra: ‘O’Reilly Media.

van Inghelandt, Delphine; Melchinger, Albrecht E., Lebreton, Claude; Stich, Benjamin (2010): Population structure and genetic diversity in a commercial maize breeding program assessed with SSR and SNP markers. In Theor Appl Genet 120 (7), pp. 1289–1299. DOI: 10.1007/s00122-009-1256-2.

Venables, William N., Ripley, Brian D. (2003): Modern applied statistics with S. 4. ed., corr. print. New York [u.a.): Springer (Statistics and computing).

Wickham, Hadley (2016): ggplot2. Elegant Graphics for Data Analysis. With assistance of Carson Sievert. 2nd ed. 2016. Cham: Springer International Publishing (Use R!). Available online at https://ebookcentral.proquest.com/lib/kxp/detail.action?docID=4546676.

Zavinon, Fiacre; Adoukonou-Sagbadja, Hubert; Keilwagen, Jens; Lehnert, Heike; Ordon, Frank; Perovic, Dragan (2020): Genetic diversity and population structure in Beninese pigeon pea [Cajanus cajan (L.) Huth] landraces collection revealed by SSR and genome wide SNP markers. In Genet Resour Crop Evol 67 (1), pp. 191–208. DOI: 10.1007/s10722-019-00864-9.

Zeng, Weishan; Su, Yan; Huang, Rong; Hu, Dehuo; Huang, Shaowei; Zheng, Huiquan (2023a): Insight into the Complex Genetic Relationship of Chinese Fir (Cunninghamia lanceolata (Lamb.) Hook.) Advanced Parent Trees Based on SSR and SNP Datasets. In Forests 14 (2), p. 347. DOI: 10.3390/f14020347.

Zeng, Weishan; Su, Yan; Huang, Rong; Hu, Dehuo; Huang, Shaowei; Zheng, Huiquan (2023b): Insight into the Complex Genetic Relationship of Chinese Fir (Cunninghamia lanceolata (Lamb.) Hook.) Advanced Parent Trees Based on SSR and SNP Datasets. In Forests 14 (2), p. 347. DOI: 10.3390/f14020347.

Zhao, Yan-Jie; Hosoya, Tsuyoshi; Baral, Hans-Otto; Hosaka, Kentaro; Kakishima, Makoto (2013): Hymenoscyphus pseudoalbidus, the correct name for Lambertella albida reported from Japan. In Mycotaxon 122 (1), pp. 25–41. DOI: 10.5248/122.25.

Zheng, Xiuwen; Levine, David; Shen, Jess; Gogarten, Stephanie M., Laurie, Cathy; Weir, Bruce S. (2012): A high-performance computing toolset for relatedness and principal component analysis of SNP data. In Bioinformatics (Oxford, England) 28 (24), pp. 3326–3328. DOI: 10.1093/bioinformatics/bts606.

